# Foveated Light-Field Compound Imager

**DOI:** 10.64898/2026.03.23.713670

**Authors:** Yan Hunag, Corey Zheng, Zijun Gao, Wenhao Liu, Shu Jia

## Abstract

Artificial vision systems hold transformative potential for biomedical imaging, diagnostics, and translational research by emulating and extending the capabilities of biological eyes. However, current techniques often face intrinsic trade-offs between spatial resolution, field of view, and depth perception, particularly in compact, biologically relevant settings. Here, we introduce FOLIC, a foveated light-field compound imaging system, which integrates compound-eye-inspired wide angular coverage and chambered-eye-inspired spatial acuity within a unified multi-aperture concave architecture. FOLIC naturally generates peripheral, blend, and foveated zones from a single capture, enabling seamless, depth-extended, multiscale visualization from wide-field context down to single-cell lateral resolution. We validate FOLIC across diverse fluorescent and non-fluorescent specimens, including cellular phantoms, tissue sections, and small organisms, demonstrating its versatility and scalability for biomedical research and related translational applications. We anticipate FOLIC to offer a biologically informed design blueprint for future artificial vision systems.

**Teaser:** A bioinspired system unifies compound and chambered eye principles to achieve wide-field volumetric microscopy.

## INTRODUCTION

Biological vision systems have evolved a wide range of architectures to meet ecological demands and fulfill survival tasks, such as predator evasion, prey localization, and environmental navigation (*1-3*). Among these adaptations, two predominant types, the compound eyes of arthropods and the chambered eyes of vertebrates, exhibit fundamentally distinct structural and functional characteristics (*4*). Compound eyes comprise numerous ommatidia arranged in a convex geometry, offering panoramic vision and exceptional motion sensitivity. In contrast, chambered eyes utilize a single optical axis to project images onto a concave retinal surface, enabling high spatial resolution and depth perception. Despite their distinctions, both visual architectures represent evolutionary strategies that are optimized for distinct ecological niches, thereby stimulating the development of bio-inspired imaging technologies.

Indeed, the structural and functional features of natural vision systems have catalyzed the advancement of artificial imaging platforms that replicate and harness these biological advantages (*5*). For instance, a notable convergence in both compound and chambered eye architectures is their utilization of curved optical surfaces, which inherently expand the field of view (FOV), reduce off-axis optical aberrations, and facilitate efficient spatial sampling within lightweight and compact forms (*6*). Inspired by these advantages, curved imaging sensors have been developed to mimic the geometry of biological retinas, improving optical performance by aligning sensor surfaces with the curved focal wavefronts of lenses (*7-9*). Nonetheless, planar sensors remain prevalent in artificial vision systems due to their high pixel densities, mature fabrication infrastructure, and broad adapt-ability to diverse existing modalities.

Expanding on these concepts, artificial imaging systems have begun to incorporate specialized architectural features from biological vision to target specific performance domains. For example, compound-eye-inspired devices emphasize ultra-wide FOV (*10-12*), extended depth of field (*13*), and rapid angular sampling (*14, 15*), making them particularly advantageous for robotic navigation (*16-19*), motion tracking (*20-22*), and wide-area biomedical surveillance (*23-25*). In contrast, artificial chambered-eye systems incorporate specialized features such as foveated vision (*26, 27*), camouflage-breaking mechanisms (*28*), and enhanced spatial sampling (*29*). These properties benefit tasks requiring high precision, detailed image analysis, and long-range target recognition (*26, 30, 31*). Despite their promising capabilities, each bioinspired design retains inherent limitations of its biological counterpart: artificial compound eyes are often constrained by their low numerical aperture and resolution due to densely packed miniature lens arrays, while artificial chambered eyes typically necessitate physical rotation or complex optical components to expand their angular and off-axis coverage (*32, 33*).

Recent hybrid imaging systems have sought to merge complementary optical features, for example, by incorporating scanning prism pairs (*34*), fisheye ball lenses (*35*), or liquid-tunable optics (*36, 37*), demonstrating improvements in angular acuity, FOV, and focal tunability. However, these approaches frequently depend on complex modulation schemes, multi-camera architectures, or bulky hardware, and are predominantly tailored for macroscopic imaging tasks. Consequently, bioin-spired platforms that integrate and exploit the complementary strengths of both biological vision models—panoramic coverage from compound eyes and high spatial resolution from chambered eyes—remain significantly underexplored, presenting an opportunity for innovative advancements in artificial vision.

Here, we present FOLIC, a bioinspired foveated light-field compound imaging system, which simultaneously delivers wide-field large-FOV coverage and high-resolution 3D volumetric vision at the single-cell scale. The technique employs a multi-aperture concave architecture that integrates the modular ommatidial layout characteristics of arthropod compound eyes with the high acuity, on-axis focal properties of vertebrate concave chambered eyes. Notably, the reconstructed FOLIC images inherently contain three distinct functional zones, peripheral, blend, and foveated, each contributing uniquely to wide-field context, scalable depth perception, and high-resolution volumetric reconstruction, respectively. We validate the performance of FOLIC across a range of phantom and biological assays, including multi-layer fluorescent microsphere phantoms, hydrogel-embedded live cells, tissue sections, and small animal specimens. These results demonstrate FOLIC as a compact, scalable imager that achieves wide-field context with targeted, high-resolution volumetric detail. This integrated capability broadens the potential applicability of bioinspired imagers, particularly in biomedical research, diagnostics, and translational domains.

## RESULTS

### FOLIC concept and characterization

In contrast to existing techniques that replicate singular aspects of biological vision, FOLIC integrates a multi-aperture concave ocular architecture, inspired by both arthropod compound ommatidia and vertebrate foveae (**Fig. 1A, Table S1, Supplementary Text S1**). Owing to its concave geometry, the array reduces close-range off-axis aberrations, as indicated by optical modeling and simulations (**Supplementary Text S2, Fig. S1**), compared to traditional convex or planar configurations, and facilitates pixel utilization for spatio-angular discrimination. This hybrid design enhances spatial sampling, angular coverage, and effective resolution through substantial inter-ommatidial overlap, producing a wide FOV centered on an angle-resolved, high-acuity foveated zone (**Fig. 1B, Supplementary Text S3**). This FOV can be modeled through ray tracing and scales linearly with imaging distance (**Fig. S2A, Supplementary Text S4**). Consequently, FOLIC realizes wide-field and high-resolution volumetric reconstructed regions using light-field approaches (*38*) (**Fig. S2B, Supplementary Text S5**). Furthermore, each microlens incorporates a custom-fabricated logarithmic axicon profile that generates a quasi-nondiffracting, extended depth of focus (EDOF) while maintaining uniform lateral and axial intensity distributions (*39*) (**Supplementary Text S6**). Whereas the curved spherical lenslets experience a defocusing problem, this strategy ensures that all elemental images remain in focus, enabling seamless accommodation of the concave optical wavefront onto a planar sensor (**Fig. S2C-D, Materials and Methods**).

**Fig. 1.**
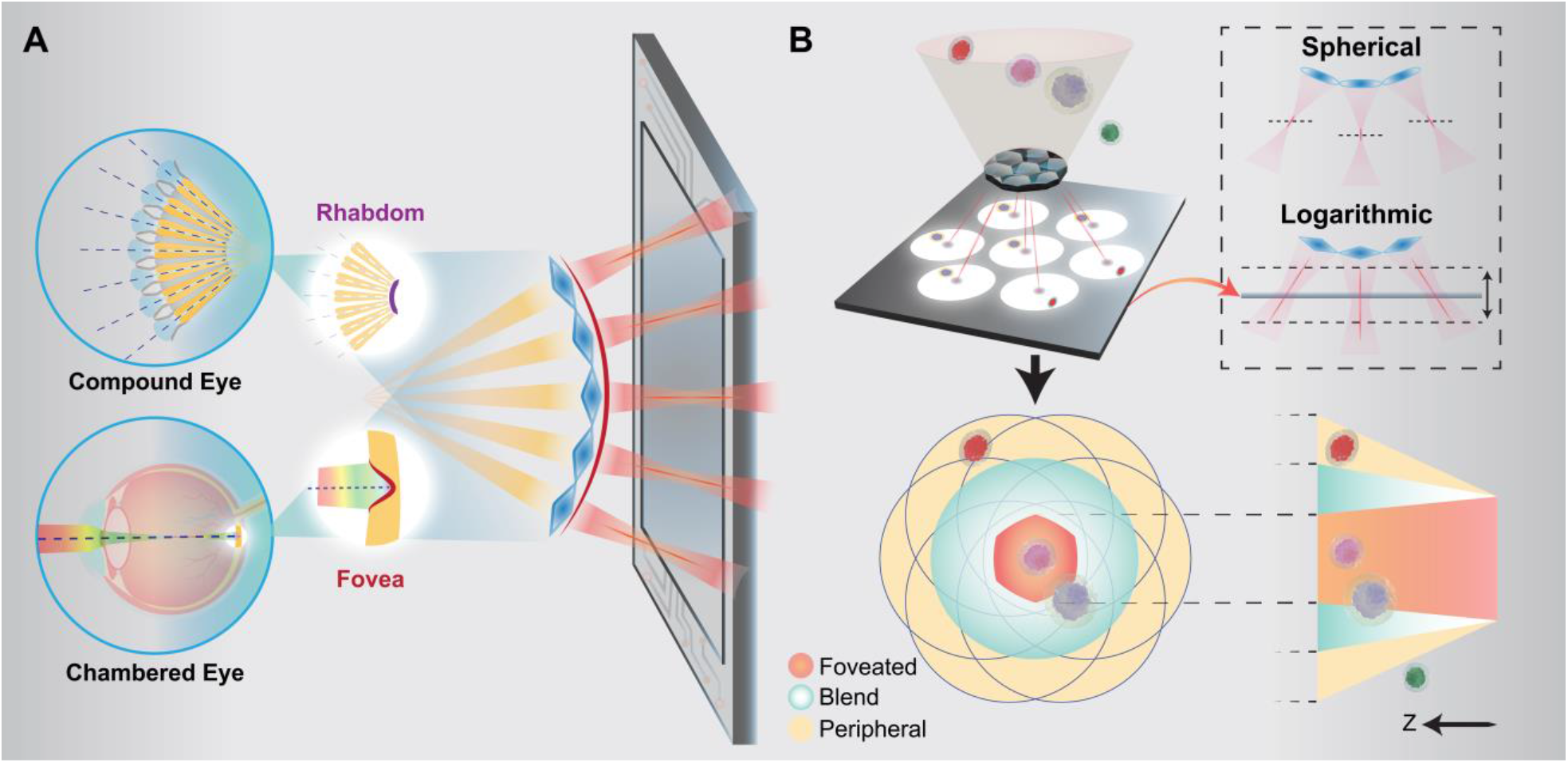
Principle of FOLIC. **(A)** Schematic overview of FOLIC, a bioinspired hybrid camera that employs a multi-aperture concave architecture that integrates the modular ommatidial layout characteristics of arthropod compound eyes with the high acuity, on-axis focal properties of vertebrate chambered eyes. **(B)** Framework of FOLIC, illustrating elemental image formation and reconstruction. The upper-right inset shows the EDOF produced by the logarithmic lenslets. The FOLIC images feature three distinct functional zones: peripheral, blend, and foveated, each uniquely contributing to wide-field context, scalable depth perception, and high-resolution volumetric reconstruction, respectively. The illustration was generated using Adobe Illustrator and Autodesk Fusion.

FOLIC was constructed from off-the-shelf components and 3D printing, with a 0**°** emission filter to block the illumination light and a simple aperture to constrain image crosstalk (**Fig. 2A, Materials and Methods**). The design has a compact form factor of 2.78 cm × 3.72 cm × 0.6 cm and can be reduced to the sensor’s lateral footprint with a cropped emission filter (**Fig. 2B, S3A-B**). The array was arranged in a hexagonal pattern (100% fill factor) and precisely tilted to optimize photon collection, magnification, and resolution (**Materials and Methods**). Specifically, the hexagonal compound arrangement was employed to facilitate radial symmetry and enhance photon efficiency with a minimal yet scalable number of lenslets. The concavity of the compound architecture was determined by the desired focal intersection geometry equivalent for each lenslet (intersection radius *R* = 3 mm; *f* = 1,500 µm and 1,750 µm for the central and edge lenslets, respectively, compensating for fabrication deviation seen through **Figure S3C-H**). The design ensures that the optical axes of all lenslets converge onto a mutual imaging zone, thereby achieving symmetric angular sampling and enhanced volumetric resolution within the foveated region. Furthermore, the logarithmic surface and intensity profile closely matched the theory (**Fig. 2C**) and extended the focal range compared to spherical lenses, allowing for all lenslets to remain in focus on the planar sensor under illumination, consistent with the numerical simulation (**Fig. 2D**, experimental setup in **Fig. S4**). FOLIC demonstrated a lateral resolving power of 4-11 µm (**Fig. 2E**) within the primary imaging zones (**Fig. S5**), matching the minimum spot size measured in the PSF (**Fig. 2F**), spanning a nearly 2-mm axial detection range (**Fig. S6**). Similarly, the axial resolution increases as the imaging distance decreases, ranging from 60 to 180 µm from the reconstructed PSF (**Fig. 2G, S7 to S9, Supplementary Text S7-8**). Notably, FOLIC maintains a high acuity axial imaging range of approximately 1.6 mm, exceeding 6-fold improvement compared to spherical lenslets (**Fig.2H, S6, S10**). These performance metrics are consistent with the resolution and volumetric requirements of cellular and tissue-scale imaging, positioning FOLIC competitively within the current state of the art (**Table S1**). While the present design integrates only a miniature microlens array (1.1 × 1.1 × 0.1 mm^3^) as the optical imaging component, the underlying design concept can be adapted for broader imaging applications (**Supplementary Text S9, Fig. S11**).

**Fig. 2.**
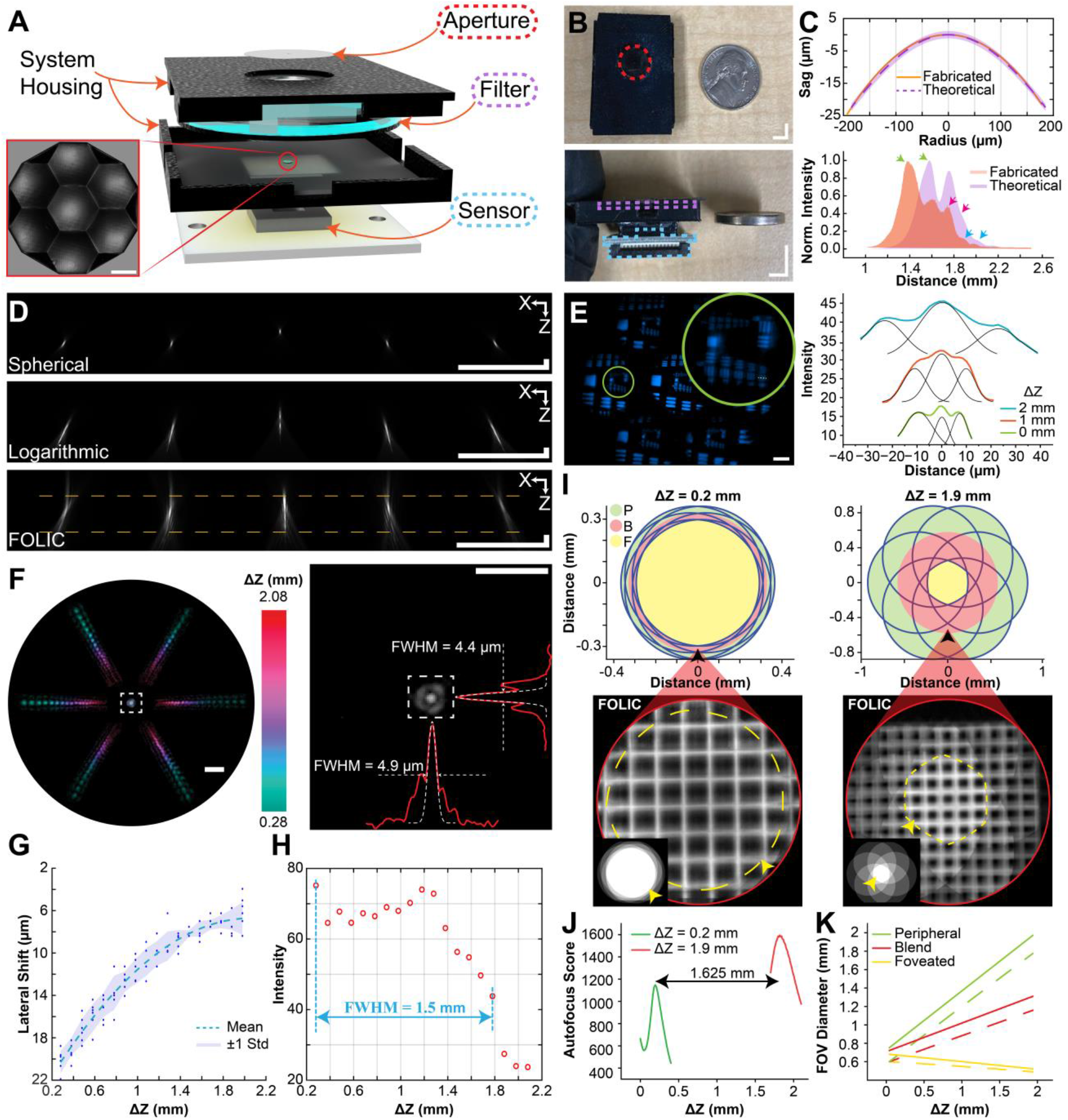
Characterization of the FOLIC system. **(A)** Illustration of the FOLIC device. The inset shows a confocal surface scan of the fabricated concave lens array. **(B)** Top and side views of the assembled FOLIC device, with dimensions highlighted relative to a U.S. quarter coin. Components are color-coded consistently with (A). **(C)** Top: Comparison of measured and theoretical surface profiles of the central logarithmic microlens. Bottom: Experimental intensity profile versus theoretical predictions under plane-wave illumination, where arrows indicate a constant offset. **(D)** Top: numerical simulation of beam propagation through spherical and logarithmic lens arrays under plane-wave illumination. Bottom: experimental measurement of FOLIC beam propagation under 635-nm plane-wave illumination. All images represent projections of five of the seven elemental images. **(E)** Experimental assessments of spatial resolution, demonstrating a minimum resolvable feature size of 4.38 µm (group 6, element 6, USAF 1951 resolution target). **(F)** Axial stack projection of the point-spread function (PSF) within an axial range from 0.28 mm to 2.08 mm, cropped to fit on a circular canvas (left). The right panel shows the PSF element of the central lenslet. (**G**) Corresponding on-sensor lateral displacement as a function of the axial position. (**H**) Intensity projection profile of the PSF element of a side lenslet. The scale bar is marked for the sensor plane. **(I)** Theoretical (top panels) and experimental (bottom panels) characterization of the field of view (FOV) at Δ*Z* = 0.2 mm and Δ*Z* = 1.9 mm. The bottom panels show reconstructed images within the blend and foveated zones. Insets show the entire FOV. **(J)** Autofocus-based depth estimation matches theoretical axial translations within 5% error. (**K**) Comparison of theoretical predictions (dashed lines) with experimental measurements (solid lines). Scale bars: 5 mm (B), 50 µm (E), 200 µm (A, D, G).

### Image formation and reconstruction of FOLIC

The elemental images acquired by FOLIC underwent an end-to-end processing framework, including aperture-based segmentation, depth-dependent lateral shifting, intensity shading correction, artifact suppression via feature mapping, and depth-dependent magnification scaling (**Materials and Methods, Fig. S12, Supplementary Software**). Notably, the reconstructed FOLIC images organize the imaging field into three functionally distinct zones, peripheral, blend, and foveated, each contributing uniquely to comprehensive visual capabilities (**Table S2**). Specifically, the *peripheral region*, viewed by one or a few lenslets, provides wide-area 2D coverage extending up to 2 mm in diameter, exhibiting minimal angular diversity and thus offering primarily spatial context without volumetric retrieval capability. Surrounding the central field, the *blend region* features inter-lenslet overlap, enabling moderate angular parallax that supports basic 3D reconstruction across an intermediate spatial zone with an average diameter of approximately 1 mm. At its core, the *foveated region* benefits from the dense convergence of all lenslets, yielding high angular sampling and high-resolution volumetric reconstruction with enhanced depth accuracy and image contrast, over an average diameter of approximately 600 µm (**Table S3**). Collectively, these three zones allow FOLIC to deliver seamless integration of wide 2D coverage with scalable depth perception and targeted high-resolution 3D visualization, all within a unified compact imaging platform. The theoretical prediction (**Fig. 2I, top**) corroborates with the reconstructed FOV (**Fig. 2I, bottom**) at two different depths, where the relevant shift estimation (1.625 mm) matches the expected sample stage translation (1.7 mm) (**Fig. 2J**). The full spatial FOV plot is shown in **Figure 2K** (maximum angular FOV in **Fig. S13**).

### Volumetric microscopy with extended depth and single-cell resolution

Conventional bioinspired imagers, including compound-eye and chambered-eye architectures, typically encounter inherent trade-offs between spatial resolution, depth perception, and FOV. Notably, the potential for volumetric retrieval remains largely unexploited in previous implementations, limiting overall performance (*40-42*). In addition, current artificial techniques have predominantly focused on macroscopic domains such as autonomous navigation or environmental surveillance, leaving microscopic and mesoscopic imaging, particularly at cellular and tissue scales, relatively underexplored (**Table S1**).

Here, FOLIC addresses these limitations through a hybrid design that leverages mutual imaging zones to enable both high-resolution spatial sampling and angular sensitivity while achieving a wide FOV through the concave multi-aperture optical structure. To validate this capability, we first captured 15-µm fluorescent microspheres distributed in the 3D space and compared the results with scanning wide-field images (**Fig. 3A-C**). As seen, the elemental images of FOLIC captured both spatial and angular information, allowing for the rendering of peripheral, blend, and foveated zones from a single light-field capture without compromising spatial details (**Fig. 3D-F**). Here, the peripheral zone exhibited a diameter of approximately 1.27 mm, and the blend zone provided volumetric reconstruction within a region measuring 0.89 mm × 0.89 mm × 0.6 mm. Importantly, the foveated zone, extending approximately 0.61 mm × 0.61 mm × 0.6 mm, demonstrated enhanced lateral and axial resolution by revealing the phantom structures of 13.84 µm (**Fig. 3F**) and 119 µm (**Fig. S14**), respectively. These results closely matched the characterized FOV (**Fig. 2K**) and EDOF (**Fig. S6, S10**) and corresponding wide-field scanning images (**Fig. 3G-I**).

**Fig. 3.**
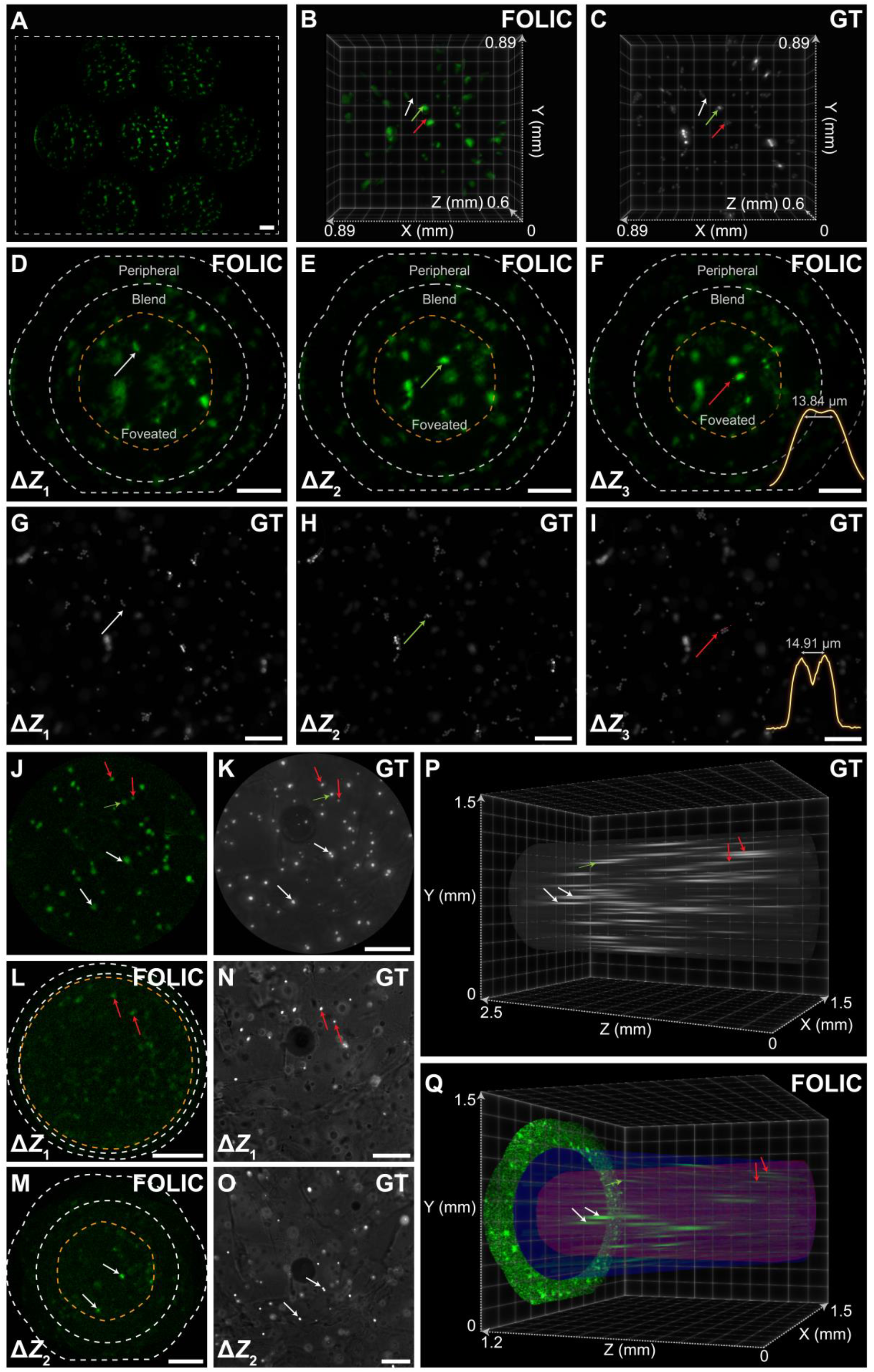
FOLIC with extended depth and single-cell resolution. **(A-C)** Elemental (A), reconstructed FOLIC (B), and wide-field ground-truth (C) images of 15-µm fluorescent microspheres distributed in three layers. The dashed box in (A) outlines the sensor area. The arrows point to exemplary consistent features. **(D-F)** Reconstructed focal images by FOLIC at Δ*Z*_*1*_ = 708 µm, Δ*Z*_*2*_= 837 µm, and Δ*Z*_*3*_= 963 µm, where peripheral, blend, and foveated zones were outlined. **(G-I)** Corresponding wide-field images at each layer, respectively. The arrows point to exemplary consistent features. Microsphere structures as narrow as ∼14 µm were consistently resolved in (F) and (I). **(J)** Elemental image of the central lenslet, which was employed as a mask to suppress noise and reconstruction artifacts. The green arrow indicates a pre-scaling feature displacement. **(K)** Corresponding wide-field images projected with 1250 axial stacks (step size = 2 µm). **(L, M)** Focal images of live HeLa cells embedded within a 2-mm-thick hydrogel, reconstructed by FOLIC at Δ*Z*_*1*_ = 192 µm, Δ*Z*_*2*_ = 966 µm. Peripheral, blend, and foveated zones were outlined. **(N, O)** Corresponding wide-field images at each layer, respectively. The arrows point to exemplary consistent features. **(P, Q)** Wide-field volume (P) and the corresponding reconstructed volume captured by FOLIC (Q), with clearly indicated foveated (red cones), blend (blue cones), and peripheral zones (single-layer view). Scale bars: 200 µm.

To further demonstrate its applicability in biological scenarios, FOLIC was used to image live HeLa cells embedded within a 2-mm-thick hydrogel, simulating tissue-like optical scattering and depth heterogeneity. The system recorded the incident light field in seven elemental images, allowing the reconstruction of individual cells using a single camera frame at a volume acquisition time of approximately 100 ms (**Fig. S15**). The extended depth of FOLIC using the logarithmic axicon micro-lenses (**Fig. 3J**) facilitated the capture of cells that were out-of-focus and poorly observable using wide-field microscopy (**Fig. 3K**) due to its limited DOF (typically 40 µm for a 4×/0.13NA objective lens). The fine 3D distributions of cells, spanning a range of 1.6 mm (thus approximately 40-fold enhancement; **Supplementary Text S10**), can be detected throughout the synthesized focal stacks (**Fig. 3L-M**), consistent with the measurements taken by scanning wide-field microscopy (**Fig. 3N-O**). The cross-sectional profiles revealed a variety of cell sizes, ranging from 12 to 17 µm in the FOLIC images over the reconstructed depth. Notably, the enhanced 3D reconstruction capability within the foveated region substantially extends the resolution compared to the surrounding blend region, collectively offering millimeter-scale 3D visualization. This capability is further complemented by wide two-dimensional contextual information provided by the peripheral zone (**Fig. 3P, Q**). These results demonstrated the ability of FOLIC to deliver accurate, high-resolution volumetric images over an extended depth, significantly advancing microscopic and cellular-level applications of artificial imagers.

### Wide-field and 3D foveated imaging of tissue samples and small organisms

Compared to standard microscopes, artificial imagers hold significant promise for compact, light-weight, and field-deployable microscopy. For example, investigating multicellular structures within tissue samples is crucial for biomedical research, yet it remains relatively underexplored by existing artificial platforms. Employing FOLIC, we acquired cryostat tissue sections of the mouse kidney stained with Alexa Fluor 488-conjugated wheat germ agglutinin (WGA), which binds to N-acetyl-glucosamine and sialic acid residues. Here, FOLIC captured perspective views of 15-μm-thick tissue specimens spanning a millimeter-scale FOV(**Fig. 4A**). The reconstructed image (**Fig. 4B**) simultaneously provided wide-field context and high-resolution foveated visualization, clearly delineating renal cellular morphologies and glomerular structures consistent with a conventional wide-field image (**Fig. 4C**), with spatial details finer than 10 µm (**Fig. 4D-G**). The results underscore the utility of FOLIC for hybrid high-throughput screening and targeted examination in diverse biological applications, such as imaging thick-dish cell cultures and mouse brain slices (**Fig. S16**).

**Fig. 4.**
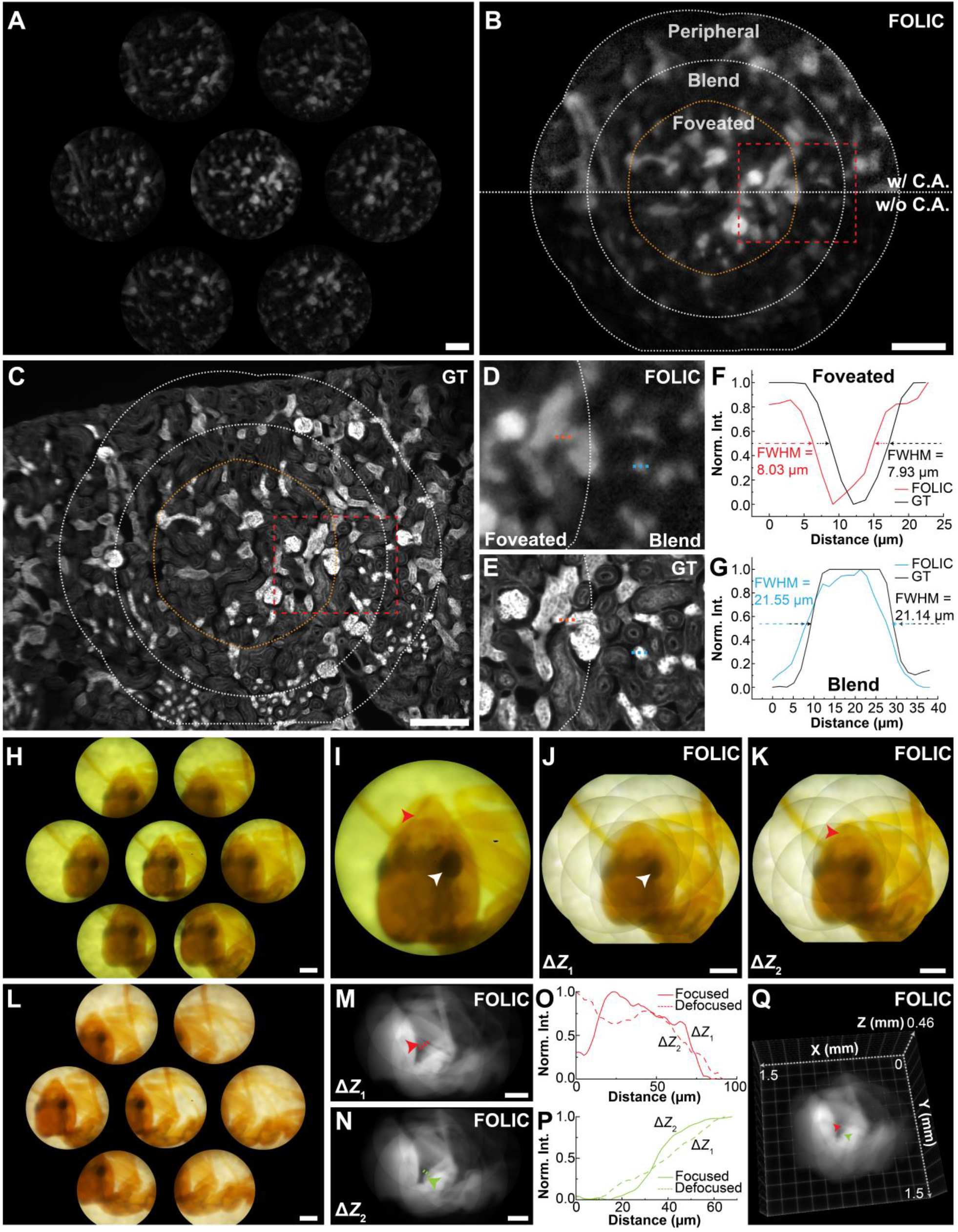
Wide-field and 3D foveated imaging of tissue sections and insect specimens. (**A**-**C**) Elemental (A), reconstructed FOLIC (B), and wide-field (C) images of a 15-μm-thick mouse kidney section labeled with WGA, featuring glycoprotein-rich regions (e.g., distal tubules). Peripheral, blend, and foveated zones were delineated. The effect of contrast adjustment was shown relative to the raw reconstruction in (B). (**D, E**) Zoomed-in images of the regions in (B, C) as marked, displaying enhanced clarity and resolution in the foveated regions compared to the blend regions. (**F, G**) Cross-sectional profiles along the dashed lines as marked in (D, E). (**H**) Brightfield elemental images of *Hymenoptera* ant specimens using fiber-based white-light illumination. (**I**) Zoomed-in image of the elemental image of the central lenslet in (H). (**J, K**) Reconstructed FOLIC images at Δ*Z*_1_= 0 µm and Δ*Z*_2_ = 111 µm with contrast adjustment. Arrows point to features in focus. (**L**) Brightfield elemental images of *Hymenoptera* ant samples using cell-phone flashlight illumination. (**M, N**) Reconstructed focal images by FOLIC at Δ*Z*_1_ = 0 µm and Δ*Z*_2_ = 168 µm, respectively. (**O, P**) Cross-sectional profiles of the structures as marked in (M, N) at the two focal planes. (**Q**) Volumetric view of the reconstructed ant specimen containing all focal layers under ambient illumination. Scale bars: 200 µm.

To further illustrate the versatility of FOLIC for studying non-fluorescent specimens, we acquired brightfield images of *Hymenoptera* ant samples. Under fiber-based white-light illumination, FOLIC captured full-color elemental images using an integrated RGB camera (**Fig. 4H-I**). The volumetric reconstruction enabled foveated synthesis of key anatomical features, including eyes and mandibles, across multiple focal planes within a depth range of approximately 170 µm and a peripheral FOV spanning up to 1.6 mm (**Fig. 4J-K**). Demonstrating practical adaptability, we further imaged specimens illuminated by a conventional cell phone flashlight (**Fig. 4L**). Even under these simplified illumination conditions, FOLIC reliably resolved microscopic structures, such as ant limb segments approximately 42 µm thick, which were clearly visualized across different focal depths (**Fig. 4M-P**) and the reconstructed volume (**Fig. 4Q**). Collectively, these results verified the robustness and adaptability of FOLIC to deliver high-quality volumetric reconstructions and precise sectioning for both fluorescent and non-fluorescent biological specimens.

## Discussion

Current artificial vision systems face inherent trade-offs among spatial resolution, FOV, and depth perception, particularly in compact and biologically relevant settings. These limitations constrain the development of integrated imagers capable of capturing both wide-area context and high-detail volumetric information simultaneously. In this study, we present the compact, bioinspired foveated light-field compound imaging system (FOLIC) that combines panoramic 2D vision with targeted high-resolution 3D volumetric reconstruction. FOLIC effectively integrates compound-eye-like angular coverage and chambered-eye-like spatial acuity within a multi-aperture concave design. The natural formation of peripheral, blend, and foveated zones within a single capture enables seamless multiscale imaging across large depths with single-cell resolution, as validated across various fluorescent and non-fluorescent biological targets under both laboratory and ambient lighting conditions. The system exhibits reduced axial resolution owing to its predominantly orthographic viewing geometry. While this is adequate for specimens with limited depth complexity or for applications where axial morphology is not the primary focus, axial performance can be further enhanced through wave-optics-based modeling or deep learning-driven volumetric reconstruction approaches (*38, 43*). Furthermore, the FOLIC framework is inherently scalable and can be further enhanced through integration with meta-optics (*44-46*), flexible optoelectronics (*47, 48*), in-sensor computing (*49-51*), camera architectures (*52-54*), spectral multiplexing (*55*), miniaturized modalities (*56, 57*), and advanced computational frameworks (*58-60*). Moreover, the underlying principle of FOLIC can be broadly adapted to augment a broad class of bioinspired artificial vision systems (*15-17, 19, 61, 62*). Collectively, these attributes position FOLIC as a versatile and expandable platform for biomedical research and selected translational applications, with potential uses in tissue mapping, cellular profiling, and field-deployable diagnostics, while offering a biologically informed design blueprint for future artificial vision systems.

## Materials and Methods

### Logarithmic lens design, fabrication, and assessment

The logarithmic lenslets were designed with a large aperture (370 µm) to enhance light collection and resolution. The central lenslet (focal range: 1,250-1,750 µm) was positioned 3mm above the sensor surface. Aligned parallel to the sensor, this lenslet captures on-axis maximum intensity projection within the detection range, providing high acuity local reference for feature mapping. To compensate for optical path length differences and fabrication deviations, the peripheral lenses were fabricated with a shifted focal range (1,500-2,000 µm) to maximize focal overlap at the sensor plane. Each peripheral lenslet was tilted at an angle of 7.057° such that the optical axes intersect 3 mm above the lenslets, corresponding to the midpoint of the detection range.

The concave lenslets were fabricated using a two-photon polymerization 3D printer (Photonic Professional GT2, NanoScribe), offering high design flexibility and sub-micron printing resolution. The fused silica substrate was initially plasma-cleaned with ozone gas to improve surface adhesion. A drop of photoresist (IP-Q, NanoScribe) was then deposited on the substrate and loaded for printing. After fabrication, the structure was submerged in SU-8 for 20 minutes, followed by a 2-minute rinse in isopropyl alcohol. Surface profiles and geometric accuracy were evaluated using an Olympus LEXT 3D Material Confocal Microscope. The intensity profiles of the printed lenslets were subsequently characterized under 635 nm plane wave illumination, confirming an overlapped detection range suitable for all-focused sensor integration. Optimized fabrication parameters for the current FOLIC lenslets are summarized in **Table S4**. A summary of the current FOLIC design parameters and performance is provided in **Table S7**.

The current design took into consideration of few realistic physical limitations (**Fig. S3A**). First, the reliable printing thickness and structural stability, with a regular 3D printer, for assembly of the system housing and spacing to prevent the emission filter from being damaged. The current emission filter was designed for 0° incident wave (plane wave) and prevents the placement between the lenslet and the image sensor, where focused excitation light by the lenses penetrates through the filter. Due to those constraints, the closest practical imaging distance is limited to ∼ 2 mm. For a proof-of-concept design, the 3 mm optical axes intersection distance was selected for an average of ∼1× magnification across the imaging range, with a single tilted lenslet layer for simplicity. The numerical aperture of ∼0.105 (diameter: 370 µm, *f* = 1.75 mm) enabled the probing of cellular resolution sampling even with reduced sampling in RGB sensors. The tilt angle was limited to 7.045°, which is constrained by the physical size of the sensor, while maintaining roughly equal FOV for each of the lenslets and meeting the 3 mm imaging distance. Therefore, by selecting different system components and fabrication approaches (e.g., microfabrication of the entire system), the design could be scaled, adapted, and further improved for performance for different imaging demands (**Fig. S17, Supplementary Text S9**).

### Fabrication and assembly flow work of the FOLIC

Upon validation of the fabricated lens array, the glass substrate was coated with black spray paint around the lenslets using a pipette gun to block stray light. The sensor (OV5647, OMNIVISION) was then stripped of its original lens unit and IR filter, exposing the sensor pixels and housing. The lenslets were integrated with the image sensor using a 3D-printed black PLA housing (Adventurer 4, FlashForge). An emission filter (89402m, Chroma) was subsequently placed above the FOLIC lenslets to enable fluorescence imaging of stained samples, in addition to bright-field imaging. A housing cover with a 600 µm aperture was then fitted above the filter to prevent image overlap. It should be noted that the use of a single aperture represents a deliberate trade-off—accepting a moderate reduction in the foveated field of view (FOV) at greater imaging depths in exchange for improved simplicity and manufacturing reproducibility. This limitation could be mitigated through more complex aperture microfabrication strategies, as demonstrated in other bioinspired camera systems (*63*).

Alignment was performed using the optical setup shown in **Fig. S3**, with the sample removed during each step of component addition. Once aligned, components were fixed in place using superglue. This proof-of-concept design can be further miniaturized by trimming the optical filter and glass substrate, reducing the system to the lateral footprint of the sensor. Detailed dimensions (**Fig. S2**) and a complete item list (**Table S5**) are provided for replication.

### System Characterization

FOLIC was experimentally characterized using sub-resolution fluorescent beads and optical targets, mounted on a motorized stage (NRT150, THORLABS). Specifically, the system point spread function (PSF) was collected by translating 2.5 µm fluorescent beads (FocalCheck, fluorescence microscope test slide #3, ThermoFisher Scientific) along the z-axis within the FOLIC imaging range, using a step size of 100 µm, sufficient for ray-optics-based volumetric reconstruction. Consequently, experimental axial resolution was evaluated from the full width at half maximum (FWHM) of the x-z and y-z intensity projections of the reconstructed PSF volume. In parallel, the field of view (FOV), lateral resolution, and modulation transfer function (MTF) (**Supplementary Text S11)** were assessed at the same 100 µm step size using a 100 µm Grid, a USAF resolution target, and a Sector Star, respectively, on a test target (R1L1S1P, THORLABS). A degree of focus score was obtained using the *Focus Measure* MATLAB program (*64*). Importantly, several layers of parafilm were stacked on the backside of the optical target to scatter the illumination for FOLIC characterization. When illuminating with a collimated light source, this effectively increases the ray angles exiting in the optical target, resembling a point source, for more accurate characterization and preventing saturation.

### Image processing and 3D reconstruction

FOLIC reconstructs a 3D volume using a light-field-based ray optics approach. At each depth, the coordinates of the edge and center lens PSF are localized, and the distances to translate the edge PSF to the center lens PSF are calculated. Next, the center PSF is set as the zero point, and a linear fit is applied to each edge lens PSF based on the located coordinates across the z-depth. From this, translation distances at unsampled depths are interpolated. The translation curves for each edge lens are then used for reconstruction.

Specifically, the FOLIC reconstruction algorithm follows a five-stage pipeline. *First*, raw elemental images are background-subtracted in ImageJ using the rolling ball method to reduce background noise and artifacts. Then, the images are segmented in MATLAB with a binary aperture mask to isolate the FOV of each lenslet. The binary aperture mask was created through thresholding of the elemental image of the physical aperture, captured through a white light illumination with the emission filter removed. *Second*, each distal segmented image (excluding the center lens) is laterally shifted by a depth-dependent translation curve calculated from the displacement of the edge elemental PSF relative to the central PSF reference. Features at the same depths converge when the segmented FOV is shifted and overlapped, allowing for synthetic refocusing of features. *Third*, a shading map (binary mask) was optionally derived from the aperture image, undergoing the same segmentation and translation. Therefore, for the current FOLIC design, the shading map contains regions from value 1 to 6, and 0 outside the FOVs for no overlaps. This mask could be optionally applied to smooth or normalize the intensity gradient in the functional zones and improve visualization, especially in the peripheral region. Technically, the *foveated, blend*, and *peripheral* zones are defined by map indices of ≥ 6, 3–5, and ≤ 3, respectively (**Fig. S12**). *Fourth*, a feature map (binary mask) reduces noise and artifacts within the foveated and blend zone volume, derived from structural features in the center lens FOV through adaptive thresholding and morphological opening. Due to the ray optics scheme, artificial features emerge as elemental images from different features that collide during reconstruction. The artifacts could be mediated by removing features not identified in the central lenslet FOV, which further enhances detail and fidelity in the reconstructed volume. Furthermore, a flat-field correction example was shown to mediate boundary artifacts (**Fig. S18, Supplementary text S12**). Lastly, the image stack is rescaled layer by layer using a pre-calibrated depth-dependent magnification factor, providing an accurate volumetric representation of the scene (see details in the **Supplementary Software**). Additionally, the co-reconstructed mask can be used to separate or visualize the three functional zones, enabling independent analysis (**Fig. 3Q**). For additional information on reconstruction efficiency and robustness (**Table S6**), including image test on artificial scattering samples (**Fig. S19**), see **Supplementary text S13**. As shown in **Fig. S19**, image contrast decreases with increasing scattering thickness, with substantial degradation observed beyond approximately one layer of parafilm (∼127 µm). This behavior is consistent with wide-field optical imaging in turbid media, where multiple scattering reduces spatial coherence and image contrast. The current FOLIC implementation is best suited for low-scattering or weakly scattering samples, including monolayer cell cultures, thin histological sections, optically cleared specimens, microfluidic assays, and surface imaging applications. The present configuration is not optimized for deep imaging in highly scattering bulk tissue without additional optical sectioning strategies. Future integration with computational sectioning, structured illumination, or confocal detection approaches may further extend performance in scattering environments.

The reconstructed resolution could be further improved through a wave-optics-based deconvolution approach, which requires finely measured point-spread functions and a higher computational cost (*38*). Alternatively, deep learning algorithms—shown to enhance both resolution and reconstruction speed in light-field systems—could be integrated into FOLIC to improve image quality, albeit also with increased computational demands (*43, 65*). These future directions highlight the potential for combining FOLIC’s compact optical architecture with data-driven computational imaging frameworks to achieve higher fidelity and faster 3D reconstruction in biomedical applications.

### Experimental imaging setup

The overall experimental setup is illustrated in **Supplementary Fig. S15A**. A blue LED (M470L3, Thorlabs) was roughly collimated to provide wide-field illumination of the sample. The emitted fluorescence was collected through a fluorescence filter set (89402m, Chroma), which transmits the emission while blocking excitation light. After image acquisition with FOLIC, the same marked sample region was imaged using a custom-built wide-field microscope, inverted to maintain identical imaging geometry. The ground truth images were obtained using a 4×/0.13 NA commercial objective lens (Plan Fluor, Nikon) and a tube lens (TTL200-A, Thorlabs). Sample scanning was performed using a motorized translation stage (NRT100/M, Thorlabs) on which the specimen was mounted. Experiment-specific setup and raw elemental image result are shown in **Fig. S15**(**B-D**).

For bright-field imaging of the Hymenoptera ant specimen, the illumination source was replaced with either a white-light fiber source or a mobile phone flashlight, diffused through an A4 sheet of paper to scatter the light. This configuration provided uniform illumination of the prepared slide while preventing oversaturation of FOLIC, which operates without the fluorescence filter protection used in fluorescence experiments. **Table S8** provides an experimental summary of the main figures.

### Preparations of layered fluorescent bead slides

The three-layer fluorescent bead slide was prepared using 15.4 µm fluorescent beads (FocalCheck, ThermoFisher Scientific), diluted to a 1:20 ratio in deionized water. First, coverslips were placed on an aluminum foil and depolarized with an electric gun (30 to 60 seconds), with the surrounding cleared, for even spreading of the bead-containing solution upon contact. Then, 20 µL of the bead solution was dispensed onto the coverslips, which were gently rotated to ensure coverage. The coverslips were then dried overnight in a closed container, away from dust and light exposure. Subsequently, the coverslips were carefully flipped, one of which was pressed onto a 1 mm glass substrate with 20 µL of glycerol mounting medium. The remaining coverslips were carefully stacked onto the previous coverslip, mounted with glycerol. Finally, the stacked coverslips were sealed with clear nail oil and dried for at least 20 minutes prior to imaging.

### Preparation, staining, and imaging of hydrogel-embedded HeLa cells

200 mg sodium alginate was added to 10 mL of FluoroBrite DMEM and vortexed, resulting in a 2 wt% alginate solution. Then, 55 mg of CaCl_2_mixed with 5 mL of FluoroBrite DMEM was pushed through a filtered syringe (with 20 μm filter pore size) to obtain a clear CaCl_2_ solution (the hydrogel precursor). HeLa cell culture was detached, centrifuged in a 15 mL tube for 5 minutes at 300 relative centrifugal force (RCF). Subsequently, the supernatant was removed, and the cells were resuspended in 1 mL of FluoroBrite DMEM. 1 mL of sodium alginate solution and 500 µL of cell medium were added to a 5 mL plastic tube and mixed. The mixture was then pipetted into a 35-mm culture dish, with 1 mL of CaCl_2_ solution prewarmed to 37°C, to form the hydrogel spheres. Then, the excess CaCl_2_ solution was removed, and 1 µM of CellTracker Green CMFDA solution in FluoroBrite DMEM was prepared and added to the hydrogel dish for whole cell labeling. The hydrogel dish was incubated for 1 hour at 37°C with 5% CO_2_. After labeling, the labeling solution was removed, and the cells were washed three times with PBS. The mixture was then pipetted into a 35-mm culture dish, with 1 mL of CaCl_2_ solution prewarmed to 37°C, to form the hydrogel spheres. The slide was sealed with transparent nail oil and left to dry for 15 to 30 minutes before imaging.

## Acknowledgments

We thank Dr. Xuanwen Hua, Keyi Han, and Zhi Ling in the Laboratory for Systems Bio-photonics at Georgia Institute of Technology for their helpful advice on this work. We thank Ronghao Zhang and Dr. Alan Emanuel at Emory University for providing the mouse brain samples.

## Author contributions

Conceptualization: Y.H., S.J.

Methodology: Y.H., C.Z., Z.G., W.L., S.J.

Investigation: Y.H., C.Z.

Visualization: Y.H., C.Z.

Supervision: S.J.

Writing—original draft: Y.H.

Writing—review & editing: Y.H., S.J

## Data, Code, and Materials Availability

All data required to evaluate the conclusions of this study, as well as the materials necessary for system reproduction, are provided in the main text and/or the Supplementary Materials. This study did not generate new materials. Additional data and codes are publicly available here: https://doi.org/10.5281/zenodo.17901088. The codes have been written in MATLAB and tested with version R2022b. The most updated version of the processing codes can be found at: https://github.com/Shu-JiaLab/FOLIC. To run the codes, unzip the compressed folder and follow the instructions in the file readme.md.

